# Deep Convolutional modeling of human face selective columns reveals their role in pictorial face representation

**DOI:** 10.1101/444323

**Authors:** Shany Grossman, Guy Gaziv, Erin M Yeagle, Michal Harel, Pierre Mégevand, David M Groppe, Simon Khuvis, Jose L Herrero, Michal Irani, Ashesh D Mehta, Rafael Malach

## Abstract

Despite the massive accumulation of systems neuroscience findings, their functional meaning remains tentative, largely due to the absence of realistically performing models. The discovery that deep convolutional networks achieve human performance in realistic tasks offers fresh opportunities for such modeling. Here we show that the face-space topography of face-selective columns recorded intra-cranially in 32 patients significantly matches that of a DCNN having human-level face recognition capabilities. Three modeling aspects converge in pointing to a role of human face areas in pictorial rather than person identification: First, the match was confined to intermediate layers of the DCNN. Second, identity preserving image manipulations abolished the brain to DCNN correlation. Third, DCNN neurons matching face-column tuning displayed view-point selective receptive fields. Our results point to a “convergent evolution” of pattern similarities in biological and artificial face perception. They demonstrate DCNNs as a powerful modeling approach for deciphering the function of human cortical networks.

---

Systems neuroscience research has been one of the fastest growing fields of science in recent years, culminating in a staggering amount of detailed depictions of neuronal functional properties. However, despite this progress, two fundamental questions remain unsolved. First, we remain largely in the dark regarding the functional role of neuronal populations in performing relevant cognitive and behavioral tasks. Second, with the exception of relatively peripheral neuronal circuits (e.g. directional selectivity in the retina (Fried, Münch, & Werblin, 2002)), no realistic model is available to demonstrate how higher order neuronal tuning-curves may actually be generated. Considering the example of the ventral stream visual recognition system, despite a large body of research depicting in great detail the neuronal face selectivities in high order visual areas both in monkeys (Tsao, Freiwald, Tootell, & Livingstone, 2006) and in humans (Davidesco et al., 2013; Grill-Spector, Knouf, & Kanwisher, 2004; Grill-Spector & Weiner, 2014; Kanwisher, McDermott, & Chun, 1997), the function these face-selective populations play in face discrimination and recognition remains unknown, with various tentative hypotheses accounting for a limited set of observations (Chang & Tsao, 2017; Haxby, Hoffman, & Gobbini, 2000; Leopold, Bondar, & Giese, 2006). These limitations are not unique to visual processes; in fact, all attempts to model high level neuronal properties rely on educated guesses that can only be loosely linked to perception and behavior. The problem is largely due to the lack of models whose functional performance can achieve realistic human or animal levels (VanRullen, 2017).

This problematic situation has been transformed in the last few years with the discovery that artificial Deep Convolutional Neural Networks (DCNN) can now approach human level performance in a variety of visual tasks, for example in face recognition (Parkhi, Vedaldi, & Zisserman, 2015; Taigman, Yang, Ranzato, & Wolf, 2014). This rapidly unfolding revolution offers the field of systems neuroscience a new type of models that achieve realistic human performance in specific tasks. Indeed, a number of recent studies have provided encouraging indications for the usefulness of DCNNs in predicting visual responses along the human visual hierarchy (Güçlü & van Gerven, 2015; Khaligh-Razavi & Kriegeskorte, 2014; Yamins & DiCarlo, 2016; Yamins et al., 2014; Yildirim, Freiwald, & Tenenbaum, 2018), as well as in capturing category-selective representational geometries in visual cortex of humans and monkeys (Cichy, Khosla, Pantazis, Torralba, & Oliva, 2016; Khaligh-Razavi & Kriegeskorte, 2014; Kuzovkin et al., 2018).

Here we employed this new approach to help resolve an outstanding question concerning the function of face selective areas in the human ventral stream: are they involved in distinguishing among different pictures of faces (pictorial function) or in the identification of personal identity across the diversity of face images (recognition function)? Our results revealed a significant and consistent similarity between the face-space geometries of human cortical face-columns, measured with intra-cranial recordings in 32 patients, and that of an artificial DCNN achieving human level face recognition performance (VGG-face (Parkhi et al., 2015)). Critically, the match was specific only to intermediate layers of the DCNN hierarchy. Targeted image manipulations revealed invariance to changes in low level features (background removal, grey scale conversion) but high sensitivity to pictorial face changes that preserved the individual identity, such as view point shifts. Finally, receptive fields of individual model neurons that were found to match specific face columns displayed view-specific face fragments such as ears and eyes (Lerner, Epshtein, Ullman, & Malach, 2008). Together, these findings highlight the power of DCNNs as a productive model of neuronal function. They further argue for a pictorial rather than a recognition function of high order face-selective cortical columns in the human visual cortex.

## Results

Visual responses to three independent sets of images, including human faces and four additional categories, were recorded using either subdural or depth intracranial EEG (iEEG) electrodes (see methods for stimuli and recording details). **Figure 1A** depicts the 1-back experimental design in which patients were instructed to view images presented for 250ms (sets 1-2) or 500ms (set 3) and press a button for image repeats. Each specific image within a category (termed here “exemplar”) was presented 3-6 times in a pseudo-random order. Altogether, data from 8924 contacts from 59 patients were analyzed. We focused on the high frequency amplitude (HFA, 48-154 Hz) signal in these recordings, following previous findings showing that this signal reveals cortical column-scale functional selectivity (Privman et al., 2007; Winawer et al., 2013), and also serves as a reliable index of aggregate firing rate in humans (Manning, Jacobs, Fried, & Kahana, 2009; Mukamel et al., 2005; Nir et al., 2007) and in monkeys (Rasch, Gretton, Murayama, Maass, & Logothetis, 2008; Ray, Crone, Niebur, Franaszczuk, & Hsiao, 2008). 94 contacts from 32 patients were found to be face selective, defined as visually responsive contacts (paired t-test, pFDR<0.05) with significantly higher response to faces relative to places and relative to patterns (two wilcoxon signed rank tests per visual contact, p<0.05; see methods for detailed inclusion procedure). 56, 52 and 23 face contacts were included in set 1, 2 and 3, respectively. **Figure 1B** depicts the cortical distribution of these face selective contacts which, as can be seen, were concentrated mainly in high order ventral visual cortex. **Figure S1** shows the distributions separately for the three sets.

**Figure 1.**
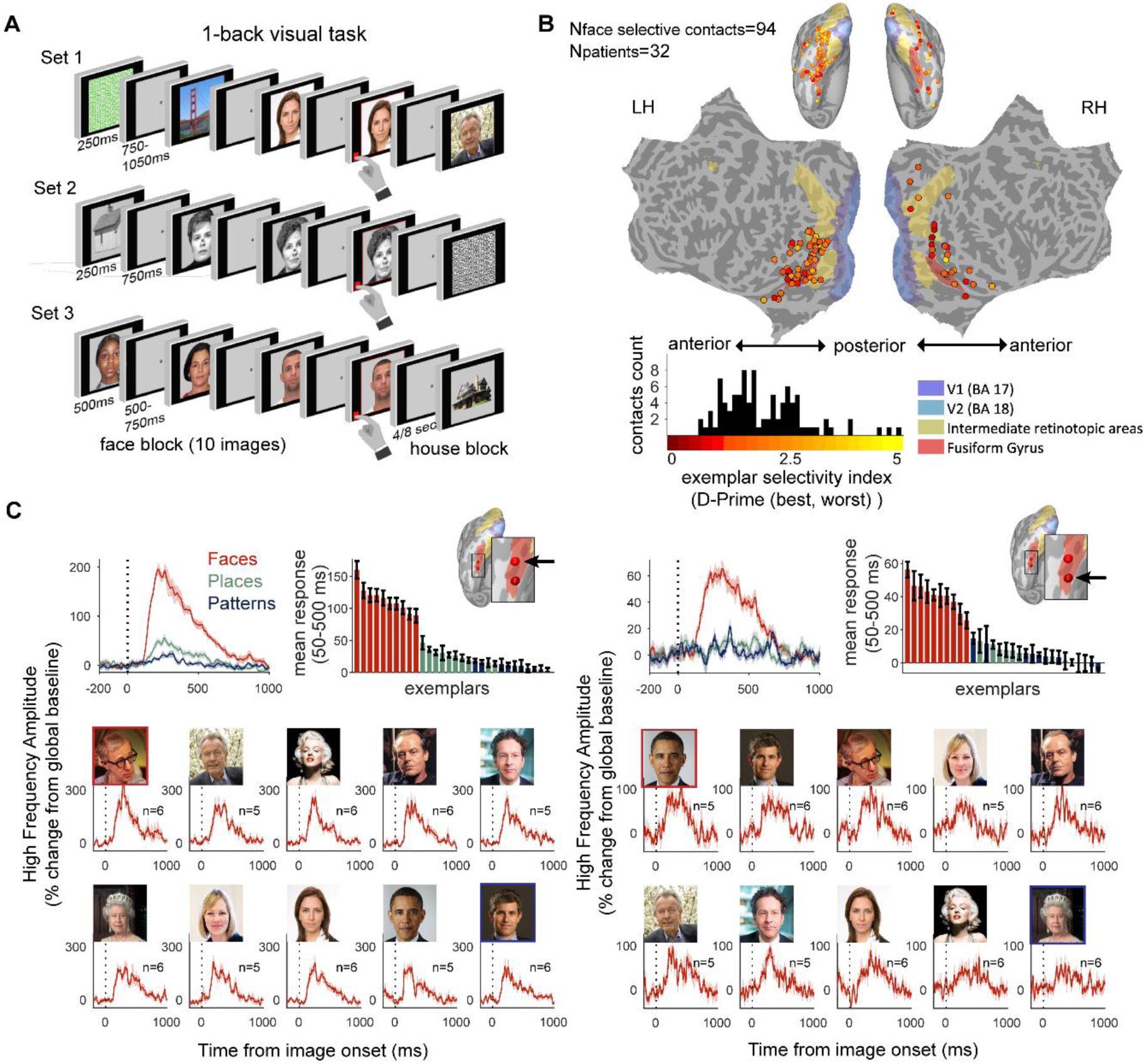
Experimental design, face selective iEEG contacts localizations and exemplar selectivity observed in individual contacts. **(A)** schematic illustration of the 1-back visual tasks. Three versions, each having a different set of face stimuli, were included and every participant took part in 1-2 of these versions. **(B)** Anatomical distribution of the face selective contacts on the inflated (ventral view, top) and flattened cortex. Color coding denotes exemplar selectivity indices, defined as the d-prime between the most preferred and the least preferred face exemplars. Histogram presents the distribution of selectivity indices alongside the color code map. The same distribution is presented separately for each of the three sets in **Figure S1**. **(C)** Example of two neighboring face contacts implanted in the same patient, their anatomical locations marked by black arrows on the inflated left hemisphere. The top left panel depicts the mean HFA response to all faces, places and patterns in set 1, demonstrating a robust face selectivity. Top right bar plot depicts the mean HFA response (50-500 ms) to each of the exemplars in descending order of activation, with bar colors denoting the visual category of the exemplar. Bottom galleries present the mean response of each contact to the different faces with the corresponding images presented on top. Galleries are presented in descending order of mean response (as in the bar plots). All error bars denote ±1SEM. HFA responses are normalized to percent signal change from the mean baseline (−200-0 ms) across all image presentations. Note the differences in response amplitude to different exemplars which was unique to each iEEG contact.

Further examination of the responses to individual face exemplars in each iEEG contact revealed substantial differences in activation amplitudes across exemplars. This can be easily discerned in the “gallery” of responses obtained from a single contact shown in the left bottom of **Figure 1C** where, for example, the response to the image of Woody Allen was more than 1.5 times greater than the response to Obama. To quantify this phenomenon, we defined an exemplar selectivity index for each face contact by computing the d-prime between its mean response to the most preferred and least preferred face exemplars. The distribution of exemplar selectivity indices and their corresponding anatomical sites are depicted by the color code mapping in **Figure 1B** and **Figure S1**. All face selective contacts showed a significant level of exemplar selectivity when compared with shuffled data (permutation test per contact, all pFDR values<0.01). Importantly, the specific profile of exemplar selectivity changed across neighboring electrodes, as illustrated in the gallery of responses of two neighboring contacts implanted in the same patient in **Figure 1C** (see also (Davidesco et al., 2013)).

At the level of contacts’ ensemble, such heterogeneous exemplar selectivity in single face contacts could potentially underlie neural discriminability between individual faces (Davidesco et al., 2013). To examine this possibility, we applied a simple pattern-matching decoding analysis. Decoded exemplar labels were assigned based on the minimal Euclidean distance between a test pattern, composed of randomly assigned single responses from each contact to each face, and a training pattern, composed of the averaged remaining responses (see methods for details). The results are presented in **Figure S2**. Indeed, decoding accuracy was significantly above chance in sets 1 and 2. Set 3 showed only a positive trend likely due to a considerably smaller number of face contacts (labels permutation test: set 1: p=0.0001; set 2: p=0.025; set3=p=0.21).

What could be the function of this face-exemplar selectivity? To examine whether a DCNN with human-level face recognition performance (VGG-Face) could serve as a realistic functional model of these selectivities we examined whether the face-space topography of face exemplars, as determined by pair-wise distances between their activation patterns (Edelman, Grill-Spector, Kushnir, & Malach, 1998; Kriegeskorte, Mur, & Bandettini, 2008), was similar between the human cortex and individual DCNN layers. Pair-wise activation distances were measured for all face exemplar pairs both in the human cortex and in each of the VGG-Face layers. For the neural data, we defined an activity pattern per face exemplar as the vector of concatenated mean responses at 50-500 ms obtained from all face-selective contacts in the relevant set. A pair-wise distance between exemplars was defined as the Euclidean distance between the two response vectors generated by a pair of faces. Comparing all the pair-wise distances generated by the iEEG recordings with those generated by the DCNN for the same face images revealed a significant correlation that was consistent across the three picture sets. Importantly, significant correlations were limited only to the intermediate DCNN layers, and chance performance was evident at early, low level feature-selective layers of the hierarchy and at the top, identity-selective layers. These results are depicted in **Figure 2**, which shows the correlation between each iEEG data set and each DCNN layer (Fig. 2 left), the scatter plots of the pair-wise distances for the two most correlated layers (Fig. 2 middle) and the actual pairs of face-exemplars presented on top of the scatter plot from the maximally correlated layer (Fig. 2 right). The correlation between iEEG recordings and two specific intermediate layers was significant in the first two sets (image permutation test followed by 0.05 FDR correction for number of layers; all pFDR values <0.05; see Fig. 2 for correlation coefficients and p values) and a similar trend was observed for two intermediate layers in the third set, which included only 23 face contacts and also showed the weakest exemplar decoding performance (non-parametric p<0.05 prior to FDR correction). A pooled analysis was performed to assess the consistency across sets. This pooled analysis is presented in **Figure 2D** and depicts the mean weighted correlations across the three sets for every DCNN layer, with set weights assigned according to the number of face contacts. As can be seen, all 5 intermediate layers, ranging from layer pool4 to pool5, significantly matched the neural distances (image permutation test followed by 0.05 FDR correction, all pFDR values<0.01).

**Figure 2.**
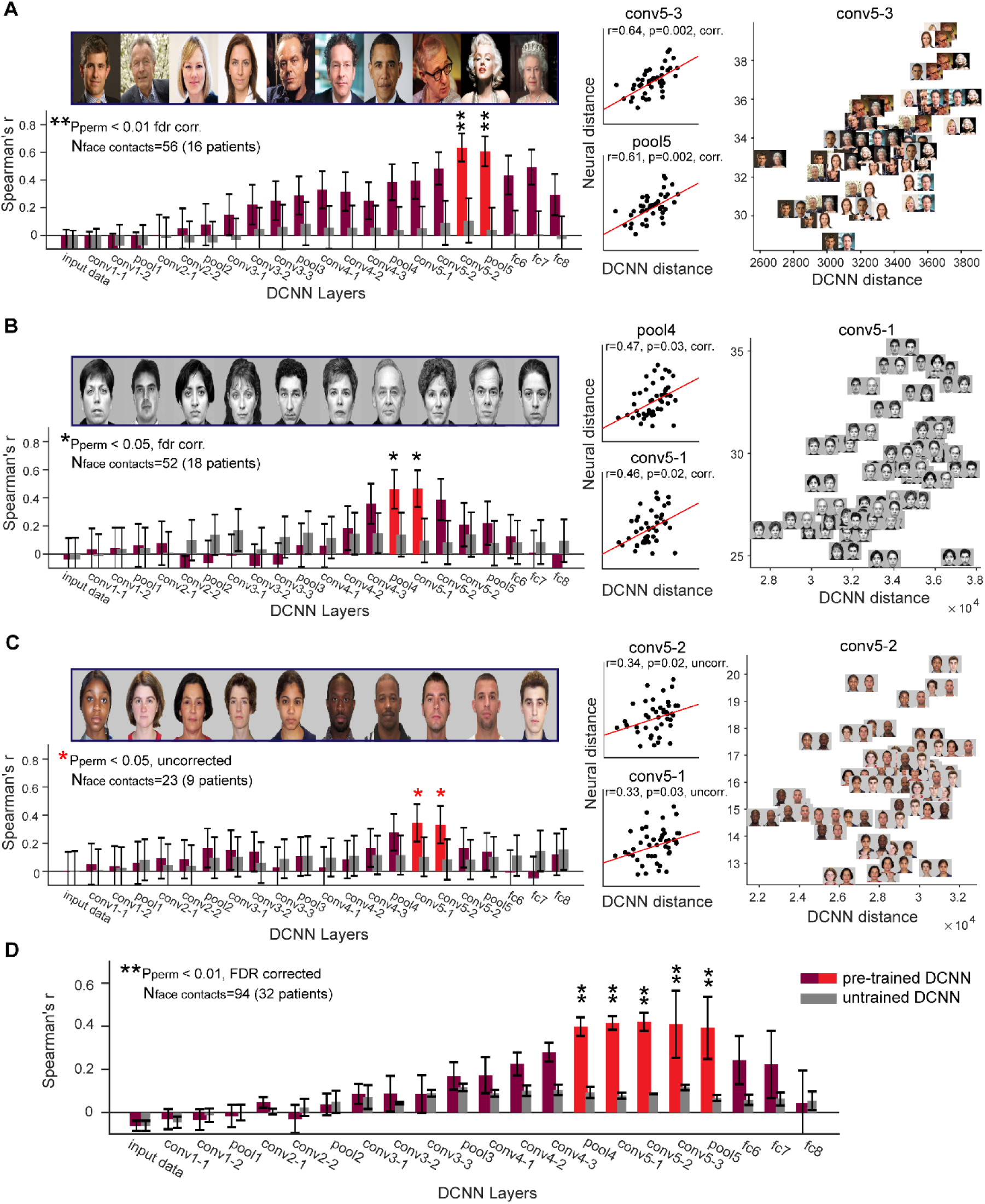
Pairwise distances in the neural face-space match the distances in intermediate DCNN layers. **(A)** Left: Purple and red bars denote the spearman’s correlation between pairwise distances in iEEG face-selective contacts and in the different layers of a DCNN pre-trained on face recognition (VGG-Face). Grey bars denote the same correlations, but to untrained VGG-Face layers. Face exemplars included in set 1 are presented above the bar plot. Error bars are the image pairs bootstrap standard error. Middle: the scatter plots depicting the correlation in the two most highly correlated DCNN layers. Red line is the least squares linear regression fit. Right: Enlarged scatter plot for the maximally correlated DCNN layer (the same data as in the top scatter plot in middle panel), with images of face pairs presented on top of individual dots. **(B)-(C)** Same as (A), only for sets 2 and 3, respectively. Note that each set consists a different group of face contacts and face exemplars. **(D)** Weighted averages of the correlation coefficients observed across the 3 sets. Sets were weighted by the number of face contacts they included. Error bars denote the weighted standard error across the sets. Note the consistent correlation of the iEEG recordings to mid-level DCNN layers. All p values were derived from a non-parametric permutation test, with 1000 random permutations of image labels followed by an FDR correction to control for a 0.05 false discovery rate across layers. Reported p values are the FDR adjusted p values, except for set3 in which significance did not survive FDR correction.

An important question is whether the effect was a result of pre-training the artificial network on a large set of face exemplars, or whether the mere deep architecture of the network was sufficient for the emergence of a match to neural distances. To examine this point, we ran the same analysis on an untrained VGG-Face network, preserving the same architecture while assigning its connections random weights drawn from a standard normal distribution. Randomizing the connection weights resulted in a near complete abolishment of the correlation to the neural face-space, as denoted by the grey bars in **Figure 2A-D** (left panel), highlighting the necessity of training the network on natural faces statistics to achieve similarity to the neural face-space.

An interesting question concerns the contribution of specific temporal dynamics in the neural responses to the observed match between neural and DCNN face-space. To assess whether such temporal dynamics hold valuable information for recapturing neural face-space through DCNN, we ran the same analysis presented in **Figure 2**, this time averaging the response across time and taking a single value – the mean HFA across the period of 50-500 ms – to denote the response of a single contact to a specific face, thus eliminating all information derived from the dynamic profile of the response. The results are presented in **Figure S2**. Despite averaging the dynamic profile, correlations in all three image sets remained significant in the same layers as prior to averaging, suggesting that the precise dynamical wave-form was not critical in generating the distance correlations to the DCNN layers.

The availability of a functional model of the neural face-space makes it possible to perform controlled manipulations on the images presented to the network and examine their impact on the match to the neural data. **Figure 3** compares the correlation to neural data given the original images as DCNN input with the correlation after applying five types of manipulations to the images presented to the DCNN. Two manipulations addressed the issue of low-level features: converting the background of the face images to black (turquoise bars), and converting the colored faces to grey-scale faces (grey bar). Both these low level manipulations had no significant effect on the brain to DCNN correlations. Conversely, presenting the same identities in a different appearance (e.g. facial expression, haircut, illumination), and presenting the same identities in similar appearance but with partial (^~^45°) head rotation – manipulations that change the appearance while maintaining personal identity – significantly reduced the correlation (pale orange bar and blue bar, respectively). Moreover, a ^~^90° rotation to a profile view completely abolished the correlation (dark blue bar). It is noteworthy that invariance to such high level manipulation can be achieved by the DCNN, but only in downstream layers to those matching the neural data: nearest neighbor classification of identities across these high-level manipulations resulted in a mean accuracy of 92% for the top fully connected layer (fc8), whereas accuracy rate for layers that matched neural data dropped to 29% and 66% (for layers conv5-3 and pool5, respectively; see **Figure S4** and methods).

**Figure 3.**
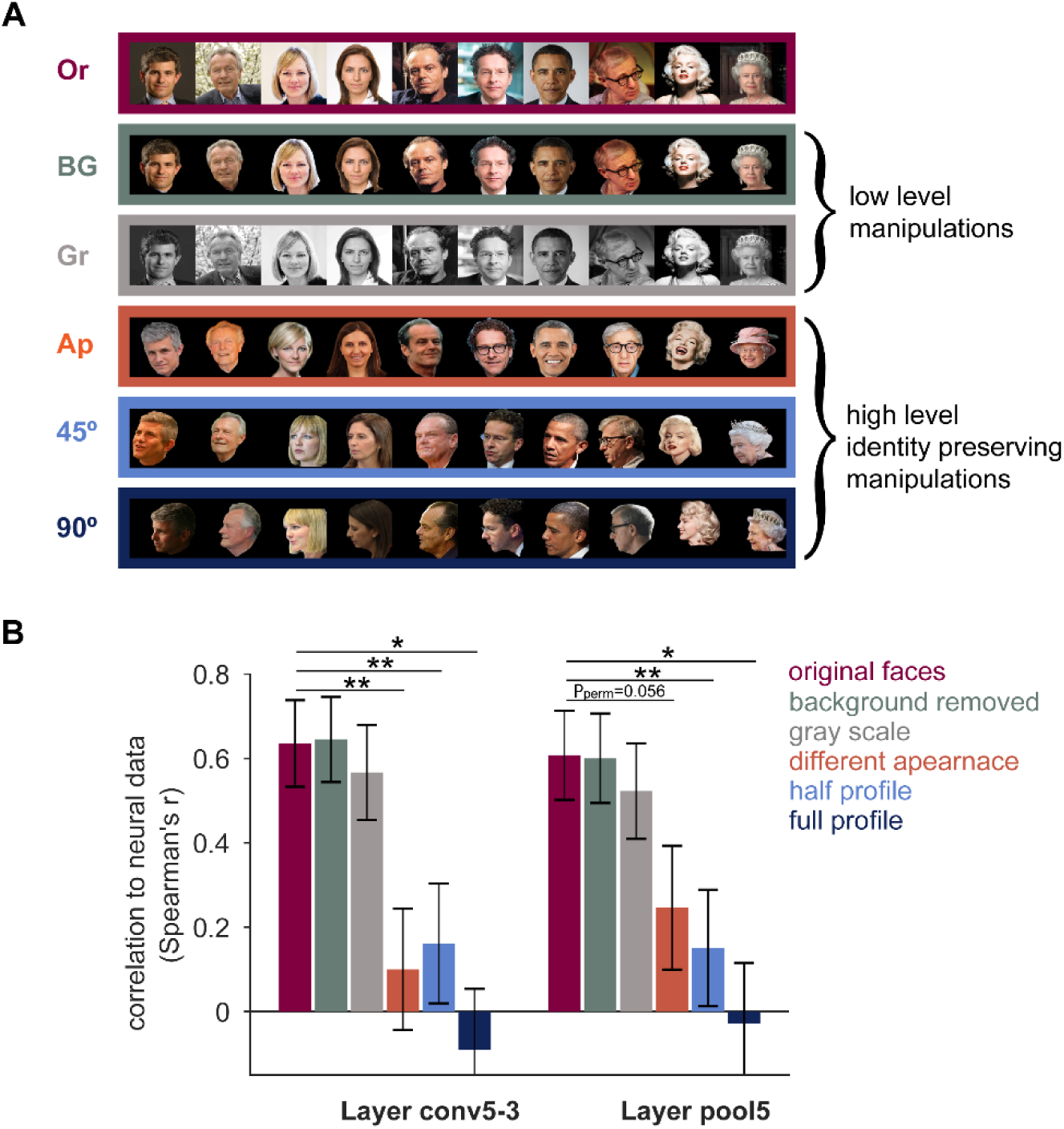
Distance similarity between neural and DCNN face representations shows invariance to low-level image manipulations but is sensitive to identity-preserving face manipulations. **(A)** Manipulated face exemplars included in set 1, from top to bottom: Or – original face images; BG – removed background; Gr – grey scale conversion; Ap – different appearance of the same identities which differ in pictorial aspects such as facial expression, illumination and haircut; 45° and 90° – different images of the same identities, rotated at ~45° and ~90° view-points, respectively. **(B)** The impact of feeding the manipulated images presented in (A) to the DCNN on its correlation to neural distances in layers that originally showed a significant correlation in set 1 (layers conv5-3 and pool5). Error bars denote bootstrap standard error on image pairs. Whereas background removal and grey scale conversion did not impact the correlation to neural data, higher level yet identity preserving manipulations (panels 3-5) significantly reduced the correlation to chance. Statistical significance was assessed by computing the same correlations 1000 times while randomly shuffling image labels in the neural data. P values were defined as the proportion of correlation reductions in the shuffled distribution that exceeded the correlation reduction obtained from the correctly labelled data. * denotes p<0.05; ** denotes p<0.01. The differential sensitivity to image manipulations points to a high-level pictorial rather than personal-identity function of the iEEG recorded face-columns.

The match of the neural face-space to specific DCNN layers opens the possibility of modelling the actual tuning properties of individual face selective iEEG contacts and exploring putative optimal “receptive field” properties that may account for their tuning properties. Note that this approach assumes a first order similarity between tuning curves of single contacts and individual artificial neurons. To examine such putative model neurons, we implemented a leave-one out search scheme in the relevant DCNN layers to search for single artificial neurons that could significantly predict the responses of specific iEEG contacts to different face exemplars (see methods for search procedure). Our results uncovered a small set of such artificial neurons, all found in set 1: two neurons from layer conv5-3 and seven neurons from layer pool5 (cluster correction applied, see methods). Every model neuron was found to significantly predict the iEEG contact responses to the held out exemplars based on a linear fit. **Figure 4** depicts two examples of such model neurons. The top left scatter plot in each panel depicts the correlation between responses to each face exemplar in the model neuron (y axis) and in a single iEEG contact recording of a neural face column (x axis), reflecting the significant similarity between the two. A particularly interesting possibility which became feasible due to the DCNN modelling is the ability to actually map the optimal stimuli, i.e. ”receptive fields”, of the model neurons. We applied two different methods for such mapping: deconvolution and activation maximization (see methods for details). Applying deconvolution on a model neuron given its response to a specific input image enables the approximate reconstruction of the fragments in the image that initially gave rise to the neuron response (Güçlü & van Gerven, 2015; Zeiler & Fergus, 2014). Activation maximization takes the approach of iteratively deconvolving (300 iterations) the receptive field and adding it to the input image, to generate an image that elicits steadily increasing responses at the model neuron. The resultant putative receptive fields appeared to share common aspects: they highlighted consistent fragments embedded in a larger part of the face image, with the right ear and left eye highlighted in the examples of panel A and B, respectively (red arrows). Nevertheless, there was a difference between the two methods, with the activation maximization revealing changes involving larger expanses of the neurons’ receptive fields.

**Figure 4.**
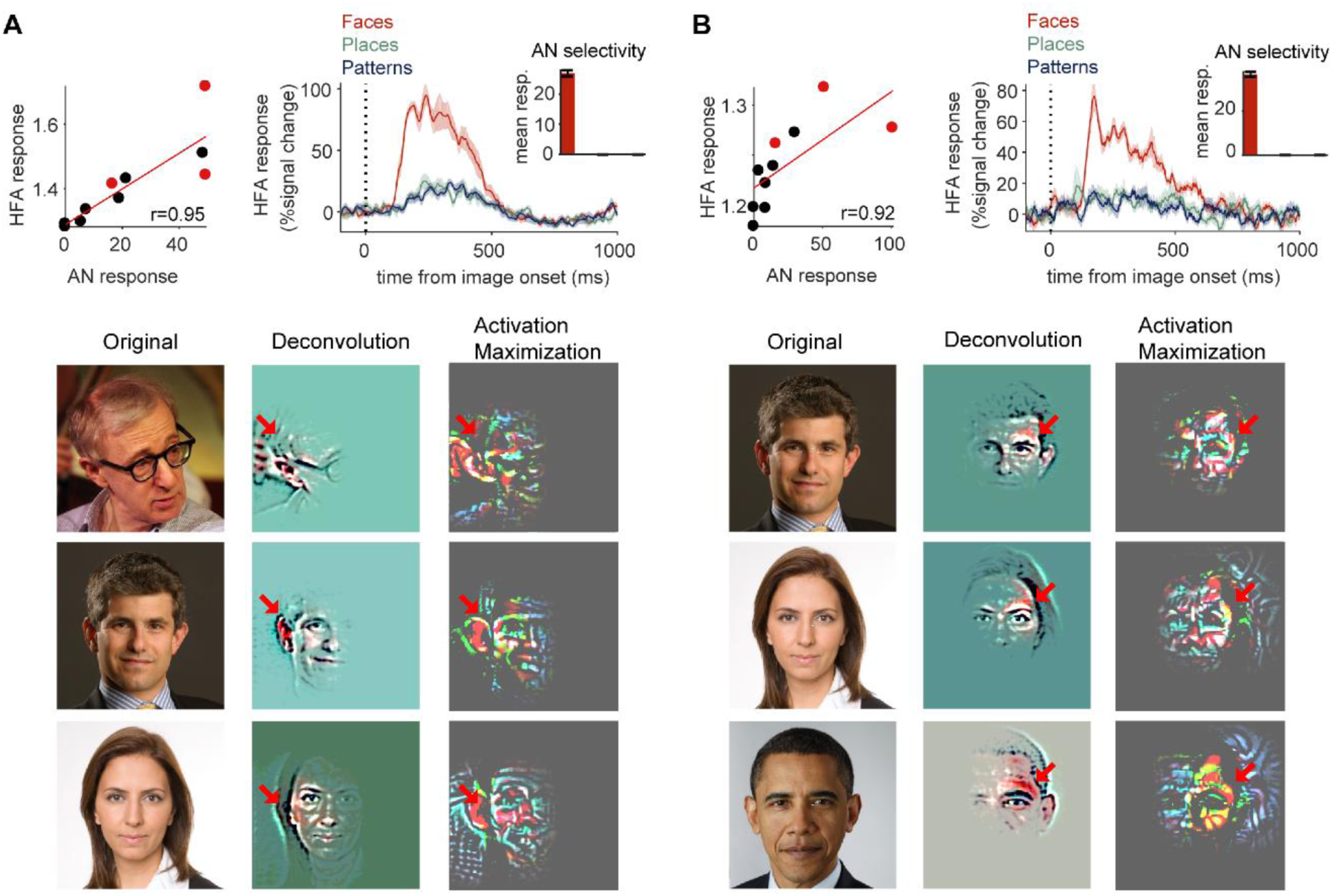
“Receptive field” reconstructions of two example model neurons. Two out of the nine model neurons that were found to significantly predict responses of single face contacts in a leave-one out search scheme (see **Figure S6** for the full set). Model neurons were defined as units which were best correlated with a single iEEG contact responses to all N-1 exemplar sets and significantly predicted the held out responses based on a linear fit. **(A)** Model neuron found in layer pool5 for set 1. Top left scatter plot depicts the responses of the model neuron (AN – artificial neuron) against the mean HFA responses (50-500ms) of the face contact. Red dots correspond to the three exemplars for which receptive field visualizations are presented (see below). Red line is the least squares linear fit. Top right plot depicts the mean categorical responses of the face-selective contact to which the model neuron matched, with shaded areas denoting standard error across exemplars. Bar inset presents the categorical selectivity of the model neuron, which responded solely to faces and not to places or patterns. Bottom gallery presents visualizations of the model neuron’s receptive field for three example face exemplars: left column shows the original images in a descending order of the model neuron’s response; middle column shows the reconstructed receptive field using the deconvolution method; right column shows the delta between the amplified image following activation maximization and the original image, reflecting the added image that maximally amplified the response of the model neuron. For visualization purpose, the contrast of each receptive field image was normalized by stretching the active range of RGB values. **(B)** Same as (A), depicting a different model neuron in layer pool5 and its matching face contact. Note the view point selectivity and consistency of the face fragments that are highlighted by the two visualization procedures, and the large expanse of image changes following activation maximization.

As can be seen in **Figure 1B** and **Figure S1**, two prominent clusters of face contacts could be discerned anatomically on the ventral surface of the left hemisphere – a posterior cluster in the inferior occipital gyrus, likely corresponding to the occipital face area (OFA) and an anterior cluster in the fusiform gyrus, likely corresponding to the fusiform face area (FFA, composed of a posterior and anterior patch in itself – pFus-faces/FFA-1 and mFus-faces/FFA-2, (Grill-Spector & Weiner, 2014)). Given the ongoing debate concerning the functional distinction between the two face patches, we separately examined their match to the DCNN layers. We defined a posterior and an anterior cluster of face contacts based on their distance to the inferior occipital gyrus and to the fusiform gyrus, respectively. We then computed the neural face-space correlation to that of the DCNN layers for each cluster separately. This was performed only for set 1 and set 2, since set 3 did not include a sufficient amount of contacts to conduct such an ROI analysis. The results of this analysis are presented in **Figure S5**. Surprisingly, for both sets, the anterior and posterior clusters independently replicated the same correlation profile across layers that we initially observed for the complete ensemble, with no significant difference in layer-selectivity arising when comparing the two clusters (permutation test on correlation difference between the two clusters, with contacts randomly assigned to clusters 1000 times; all uncorrected p values for the 5 intermediate layers > 0.46, in both set 1 and set 2).

Finally, we tested whether the observed DCNN-neural match was confined to the HFA signal or whether it could be revealed also in the iEEG evoked responses. To this end, we re-computed the neural match to DCNN layers, this time extracting the mean evoked response potential from the raw (common-referenced) iEEG signal. The analysis failed to reveal a significant correlation to any of the DCNN layers, in any of the three sets. Similarly, trial by trial extraction of the instantaneous band limited power at 8-13Hz failed to reveal a significant correlation with the DCNN face-space in any of the layers or sets.

## Discussion

The present findings reveal a significant match between the face-space of human face-selective columns recorded intra-cranially and a DCNN (VGG-Face) capable of human-level face recognition performance. The match was specific to the face pre-trained DCNN and was absent for the same network when untrained and assigned with random connection weights. These results point to an intriguing “convergent evolution” emerging between biological and artificial networks, likely driven by the similarity in task demands (Yamins & DiCarlo, 2016; Yamins et al., 2014). Understanding the parameters underlying this convergent evolution is highly informative to understanding how these networks accomplish their visual perception. This can be illustrated by considering the example of biological and artificial flying, where the common emergence of wings highlights their functional importance to the task of flying. Here, the cortical and DCNN convergence highlights the functional importance of pattern similarity to face perception and thus extends earlier theoretical proposals by Edelman and Grill-Spector (Edelman et al., 1998) of the fundamental role of such similarities in object representations as well as more recent ones in the domain of object categories (Khaligh-Razavi & Kriegeskorte, 2014; Kriegeskorte, Mur, Ruff, et al., 2008). These results are also compatible with our previous study, showing significant correlation between face exemplar activation distance measures and their perceptual similarity (Davidesco et al., 2013) (but see (Rajalingham et al., 2018)).

However, it is important to emphasize that face perception is a multi-faceted process that involves, on the one hand, a pictorial function, i.e. our essentially limitless ability to distinguish among different images of faces and, on the other hand, a recognition function, in which we can identify specific personal identities across a diverse set of different view-points and appearances. At present, it is not known whether the function of human face-selective cortical columns is mainly in the pictorial or in the recognition domain. Three converging lines of results in the present study point to a pictorial rather than personal identity recognition function of human face selective columns. First, the face-space topography match was consistently restricted to the mid-hierarchical layers of the DCNN, arguing both against strictly low-level feature representations at the bottom end and appearance-invariant recognition level at the high end (**Figure 2**). This could not be attributed to a posterior cortical bias of the recording sites since separately computing the neural-DCNN match for posterior and anterior recording sites failed to reveal a higher level match in the more anterior cluster (**Figure S5**). Second, image manipulations targeting low level features, such as removing all background information or converting the face colours to grey, failed to affect the neural to DCNN match (**Figure 3**). In contrast, higher level manipulations of the pictorial aspects of the faces which preserved their personal identity (**Figure 3**) revealed a significant reduction in the neural to DCNN match, demonstrating the dependency of the effect on some high-level pictorial aspects rather than on appearance- or view-invariant personal identity. Finally, visualizing the receptive fields of artificial neurons whose selectivity profiles matched that of specific intra-cranially recorded face-columns revealed consistent view-point selective face fragments such as ears and eyes. These fragment-like receptive fields are compatible with previous reports of the selective activation of face-areas to informative fragments (Lerner et al., 2008; G. Wang, Tanaka, & Tanifuji, 1996). Interestingly, the two different approaches we applied to reveal these receptive fields highlighted different levels of holistic representations, with a more a localized receptive field revealed through the deconvolution method while a more gestalt-like receptive field revealed through the activation maximization approach (see **Figure 4**). This dependency of the receptive field properties on their mapping method is reminiscent of the finding of a more localized RF in monkey infero-temporal cortex when a reductive method is applied (Lerner et al., 2008; G.Wang et al., 1996) vs. more holistic RF properties revealed when using face-template approaches (e.g. (Chang & Tsao, 2017; Hasson, Hendler, Bashat, & Malach, 2001; Leopold et al., 2006)).

Overall, our results strongly support a functional role for human face-selective columns in representing how face images look rather than in recognizing the personal identity of the face. Such identity recognition may be subserved by more downstream medial temporal lobe structures (Quiroga, Reddy, Kreiman, Koch, & Fried, 2005) or by the extended nodes of the face network in anterior temporal and frontal cortices that we failed to sample (Avidan & Behrmann, 2009; Freiwald & Tsao, 2010; Rosenthal et al., 2017).

In summary, the revolutionary discovery that artificial deep networks can achieve human level recognition performance offers, for the first time, a truly realistic model of high order visual processing. Together with the large scale iEEG recordings of human visual columns they allowed fresh insights into the functional role and mechanistic generation of high order human face representations. Employing the rapidly evolving new DCNNs may help, in the future, to resolve outstanding issues such as the functionalities of distinct cortical patches in high order visual areas and the functional role of top down and local recurrent processing in brain function (Spoerer, McClure, & Kriegeskorte, 2017).

## Materials and methods

### Participants

59 participants monitored for pre-surgical evaluation of epileptic foci were included in the study, 32 of them had face selective contacts (11 females, mean age 35 years with SD=11.6; see Table S1 for individual demographic, clinical and experimental details). All participants gave fully informed consent, including consent to publish, according to NIH guidelines, as monitored by the institutional review board at the Feinstein Institute for Medical Research, in accordance with the Declaration of Helsinki.

### Experimental procedures

Three versions of a 1-back visual task were included in this study, each consisting a different set of 10 face images and 4-5 additional visual categories with 10 images each from the following: patterns, places, objects, body parts, words and animals. Face stimuli in task one (set 1) were natural face images of familiar people collected in an internet search; face stimuli in task 2 (set 2) were taken from a data base previously reported (Allison, Puce, Spencer, & McCarthy, 1999); and face stimuli for task 3 (set 3) were taken from an open-source data base (Minear & Park, 2004). In all three versions, stimuli were squared and centrally presented, subtending a visual angle of approximately 13° in task 1 and 11° in tasks 2 and 3. In the first task, images from different categories were presented for 250 ms and were followed by a jittered inter stimulus interval ranging from 750 to 1050 ms. The task included 360 trials, 25 of which were 1-back repetitions. In the second task, images from different categories were presented for 250 ms at a fixed pace of 1Hz. The task included 200 trials, 25 of which were 1-back repeats. The third task was designed in blocks, where each block included 10 mages from the same category presented in pseudo-random order. The images were presented for 500 ms, followed by a jittered inter stimulus interval ranging from 750 to 1500 ms. Blocks were separated by 4/8 seconds. The task included 260 trials (26 blocks), 18 of which were 1-back repeats. During the tasks, participants were seated in bed in front of an LCD monitor. They were instructed to maintain fixation throughout the task and to click the mouse button whenever a consecutive repetition of the exact same image occurred. Five of the participants (one in set 1, two in set 2 and two in set 3) were instructed to press one of the two mouse buttons on each trial to indicate whether a 1-back repeat occurred. Every participant took part in 1-2 of the task versions.

### Electrodes implant and data acquisition

Recordings were conducted at North Shore University Hospital, Manhasset, NY, USA. Contacts were either subdural grids/strips placed directly on the cortical surface and/or depth electrodes/stereo-EEG. Subdural contacts were 3 mm in diameter and 1 cm spaced, whereas depth contacts were 2 mm in diameter and 5 mm spaced. The signals were referenced to a vertex screw or a subdermal electrode, filtered electronically (analog bandpass filter with half-power boundaries at 0.07 and 40% of sampling rate), sampled at a rate of either 512Hz or 500Hz and stored for offline analysis by XLTEK EMU128FS or NeuroLink IP 256 systems (Natus Medical Inc., San Carlos, CA). Electrical pulses were sent upon stimuli onsets and recorded along with the iEEG data for precise alignment of task protocol to neural activity.

### Anatomical localization of electrodes

Prior to electrode implantation, patients were scanned with a T1-weighted 1 mm isometric anatomical MRI on a 3 Tesla Signa HDx scanner (GE Healthcare, Chicago, Illinois). Following the implant, a computed tomography (CT) and a T1-weighted anatomical MRI scan on a 1.5 Tesla Signa Excite scanner (GE Healthcare) were collected to enable electrode localization. The pre-implant MRI was aligned with the post-implant MRI, and the post-implant MRI was aligned with the post-implant CT using a rigid affine transformation as implemented by FSL’s Flirt (Jenkinson & Smith, 2001). Concatenation of the two alignment transformations allowed visualization of the post-implant CT scan on top of the pre-and post-implant MRI scan. Individual contacts were then identified manually by inspection of the CT along with the post-implant MRI and were marked in each patient’s pre-implant MRI native space, using BioImage Suite (Papademetris et al., 2006). Electrode projection onto the cortical surface was performed as previously reported (Golan et al., 2016) (see also (Groppe et al., 2017) for an open source tool box with a similar pipeline). Individual patients’ cortical surface was segmented and reconstructed from the pre-implant MRI using FreeSurfer 5.3 (Dale, Fischl, & Sereno, 1999), and each electrode was allocated to the nearest vertex on the cortical surface. In order to project electrodes from all patients onto a single template, the unfolded spherical mesh of each individual was resampled into a common unfolded spherical mesh (reconstructed from FSAverage template), using SUMA (Argall, Saad, & Beauchamp, 2006). Colored labels on the cortical surface as presented in Figure 1B were derived from surface-based atlases as implemented in FreeSurfer 5.3: functional atlas of retinotopic areas (L. Wang, Mruczek, Arcaro, & Kastner, 2014) (intermediate retinotopic areas); Destrieux anatomical atlas (Destrieux, Fischl, Dale, & Halgren, 2010) (Fusiform gyrus); and Juelich histological atlas (V1 and V2 corresponding to Brodmann areas 17 and 18, respectively).

### iEEG signal preprocessing and HFA estimation

Signals that were initially recorded at a sampling rate of 512 Hz were down sampled to 500 Hz for consistency. Raw time-courses and power spectra of all channels were manually inspected for noticeable abnormal signals and other contaminations, and channels appearing as highly irregular were excluded from further analysis. Next, channels were re-referenced by subtracting the common average signal from the intact channels.

To estimate high frequency amplitude (HFA) modulations, the signal was divided into 9 frequency sub-ranges of 10 Hz width, ranging from 48 to 154 Hz. The sub-ranges did not include 59-61 Hz and 117-121 Hz in order to discard line noise. Each frequency sub-range was band-passed and the momentary amplitude in each sub-range was estimated by taking the absolute value of the filtered signal’s Hilbert transform (Fisch et al., 2009; Lachaux et al., 2005; Noy et al., 2015). Since the 1/f profile of the signal’s power spectrum results in greater contribution of lower frequencies to the overall HFA estimation, we normalized each sub-range by dividing it with its mean value, and averaged the normalized values across all 9 sub-ranges (Fisch et al., 2009). All data preprocessing and analyses were carried out using in house MATLAB codes. For filtering of frequency sub-ranges, we used original EEGLAB’s Hamming windowed FIR filter (pop_eegfiltnew function, (Delorme & Makeig, 2004)).

### Definition criteria of face selective contacts

Face selective contacts were defined as visually responsive contacts with a significant response to faces compared to places and compared to patterns: First, we tested whether the mean responses of each contact to all available stimuli were significantly greater then baseline (paired t-test on mean exemplar responses versus baseline, at 50-500 ms and -200-0 ms relative to image onset, respectively). Hit, miss and false alarm trials were excluded from all analyses. Note that in order to avoid a bias due to possibly different numbers of presentations for each of the images, we first averaged all responses to identical images and ran the test on the mean response to each image. FDR correction was then applied to the pooled p values from all 59 patients. Contacts with an FDR-adjusted p value smaller than 0.05 and a considerable effect size (Glass’ Δ) of larger than 1 were defined as visually responsive. 551 out of 8294 contacts were found to be visually responsive. Next, visual contacts that were significantly more selective to faces when contrasted to places and to patterns were defined as face contacts (two wilcoxon signed rank tests per contact, p<0.05, uncorrected). Finally, we applied anatomical constraints whereby face contacts located within V1, V2 or in frontal regions, as well as contacts localized further than 10mm from the cortical surface, were excluded from the set. This resulted in the final set of 94 face contacts from 32 patients: 56, 52 and 23 face contacts in set 1, 2 and 3, respectively. 33 contacts overlapped between sets 1 and 2, and 4 contacts overlapped between sets 1 and 3. The cortical distribution of face contacts is presented for each set separately in **Figure S1**. None of the detected face contacts were identified as located over the seizure onset zones by an epileptologist’s inspection.

### Exemplar selectivity index of individual face contacts

To assess the level of selectivity to different faces within each face-selective contact we defined a face exemplar selectivity index as the d-prime between the most preferred face and the least preferred face:

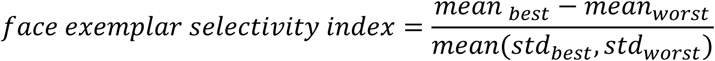

Where *mean _best_*_/*worst*_ is the mean HFA response at 50-500ms relative to face onset, and *std _best_*_/*worst*_ is the standard deviation across repetitions of the relevant face image. Selectivity indices presented in **Figure 1B** were computed on all face images available to the specific face contact. Selectivity indices presented in **Figure S1** were computed on face images included in the specific image set relevant to each of the three panels. We applied a permutation test to each face contact to assess the significance of exemplar selectivity, where the same index was computed for 1000 random shuffles of all single responses. P values were defined as the proportion of shuffle-derived indices that exceeded the original index computed on the unshuffled data.

### Decoding face exemplars from the ensemble of face contacts

To decode specific face exemplars from the activation pattern of all face-selective contacts we deployed a simple template matching decoding scheme. First, data was ordered in a three dimensional matrix where entry k,i,j, is the k^th^ trial raw HFA response (50-500 ms) to the i^th^ face exemplar in the j^th^ face contact. This matrix was generated for each of the three sets. For every set, we iteratively (1000 iterations) and randomly assigned a single trial from each face exemplar and face contact to a test pattern matrix, and averaged the remaining responses to serve as a reference pattern matrix. Thus, every decoding iteration began with a test and a reference matrix, where entry i,j is the response of the j^th^ face contact to the i^th^ face exemplar. Next, we assigned decoded labels based on the minimal Euclidean distance between the test and the reference patterns: on every matching step, the minimal distance between any test pattern and any reference pattern was detected and the corresponding label was assigned to the test pattern. The detected pair of test-reference patterns was then excluded from the subsequent match searches such that each test pattern and reference pattern could only be assigned once in every decoding iteration. The mean number of accurately decoded face exemplars across the 1000 iterations was defined as the decoding accuracy.

Statistical significance of decoding accuracy was assessed in a permutation test. The same decoding process was applied while shuffling the labels of the test patterns, resulting in a distribution of 1000 decoding accuracies generated under the null hypothesis (i.e. that single HFA responses are non-informative as to the specific face presented). P values were defined as the proportion of decoding accuracies in this null distribution exceeding the mean decoding accuracy given the original labels.

### Pair wise distances in the neural face-space

The HFA signal was smoothed with 50 ms running average window prior to computing all pairwise neural distances. For a single face exemplar, the HFA responses (50-500 ms) were averaged across repetitions in every face contact. Next, the mean responses from all face contacts were concatenated to a vector which served as the neural representation of the specific face exemplar. Pair wise distances were defined as the Euclidean distance between all representation pairs (45 distance measures per image set, identity pairs excluded).

To test the contribution of the exact wave form during the 450 ms response (**Figure S3**), we generated a neural representation based on temporally averaged responses, where the response of a single face-selective contact to an exemplar was averaged both across repetitions and across time, resulting in a single mean amplitude value. All remaining analyses addressing the match to DCNN were otherwise identical.

To investigate the specificity of the neural-DCNN correlation to the HFA signal, we computed the neural distances for two additional signals, focusing on low frequencies of the iEEG signal: the conventional ERP and the instantaneous power of low frequencies at 8-13 Hz. ERP responses were extracted by locking the common referenced raw signal to image onsets and averaging across repetitions of the same exemplar. Following (Fisch et al., 2009; Lachaux et al., 2005), low frequencies band limited power was computed by filtering the common referenced signal at 8-13Hz and extracting the absolute value of the Hilbert transformed signal. The response window for both ERP and low frequency instantaneous power was defined at 125-250 ms relative to image onset (Privman et al., 2010).

### DCNN model

We used the pre-trained VGG-Face feedforward architecture of Parkhi et al. (Parkhi et al., 2015). The network consists of five stacked blocks followed by three fully connected (fc) layers. Each block consists of 2-3 consecutive convolutional layers (13 convolutional layers in total) followed by max pooling. All 13 convolutional layers and 3 fully connected layers are followed by a rectification linear unit (ReLU), introducing non-linearity to the model. The network was trained on a large scale dataset with over 2 million face images for recognition of 2622 identities. This model reached nearly perfect performance, comparable to state of the art models (e.g. DeepFace (Taigman et al., 2014)). Importantly, none of the identities presented in our three tasks were included in the set used to train the network. Preprocessing of our raw stimulus images prior to feeding forward through the network included resizing to match network input size of 224×224 pixels and mean RGB channel value subtraction, as estimated from the network’s training set.

To compute pairwise distances in the DCNN, we passed forward the images through the network and extracted the output activations of all units at each layer. We then computed the Euclidean distance between pairs of activation patterns corresponding to image pairs, to generate a pair-wise distances vector at each layer.

### Estimating face-space similarity between DCNN layers and face-selective iEEG contacts

The neural pairwise distances vector was correlated to 22 DCNN pairwise distances vectors – corresponding to each of the VGG-Face network layers – using spearman correlation. To assess statistical significance of the correlation to each layer, we shuffled the face image labels 1000 times and recomputed the neural distances vector while leaving the DCNN distances vectors fixed. P value assigned to each layer was the proportion of correlation values derived from shuffled data that exceeded the original correlation value. An FDR correction was applied on the resultant 22 p values, to control for a 5% false discovery rate across the layers.

To estimate the consistency in correlation patterns across the three sets (**Figure 2D**), we computed the weighted average of correlation coefficients in each DCNN layer across the three sets. The weighted correlation to every layer was defined as:

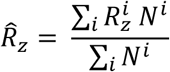

Where 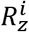 is the fisher’s z transformed spearman correlation in set *i*, and *N^i^* is the number of face contacts in set *i*. The weighted mean fisher transformed correlation was then back transformed.

### Impact of presenting manipulated images to the DCNN on the correlation to neural face-space

Low-level manipulations, including background removal and rgb to grey scale conversion of face images from set 1 (**Figure 3**, light green and grey bars), were carried out using Adobe Photoshop. For high-level face manipulations, including different appearance and two-stage view shifts, we took advantage of the faces in set 1 depicting familiar people and searched for available images of the same identities taken from a half profile and a full profile (**Figure 3**, light blue and dark blue bars), and from a similar view but under a different appearance (e.g., facial expression, haircut, illumination. **Figure 3**, light orange bar). All manipulated faces were resized and aligned to optimally overlap with their original counterparts.

To statistically test the impact of the five above mentioned manipulations on the correlation of neural data to the two DCNN layers that originally showed a significant match in set1, we applied a permutation test per manipulation and layer. In each permutation test, face labels of the neural data were shuffled 1000 times to obtain a distribution of correlation deltas – the difference between correlation given the original images as DCNN input and the correlation given the manipulated images as DCNN input. P values were then defined as the proportion of shuffled-driven correlation deltas that exceeded the real labels correlation delta.

### Decoding identity across high level manipulations from DCNN layers

To assess the DCNN capability in recognizing identity across different pictures of the same person, we pooled the images from the three high level manipulations presented in Fig. 3A (different appearance, ^~^45° rotation and ^~^90° rotation) and the original images to one data set. We then randomly assigned one image of each identity to constitute a ten images test set, and applied a Euclidean distance nearest neighbor classifier to classify the identities in the test set. We repeated this procedure 1000 times per layer, resulting in a mean classification accuracy for the two layers that matched the neural face-space in set 1 and for the top layer in the network (see **Figure S4**).

### Detection of model neurons

We attempted to find single artificial units (termed here “model neurons”) that can significantly predict the responses of single face-selective contacts to the different faces. We deployed the following search scheme per face-selective contact in each of the three sets: First, the linear correlation between the neural responses (mean HFA amplitude at 50-500 ms) and the layer’s units was computed for nine out of the ten faces. Next, we took the unit that best correlated to the 9 neural responses and used a least squares linear fit to predict the neural response to the held out 10^th^ face. We repeated this procedure ten times, holding out a different face exemplar on every iteration. If the same unit was best correlated to the neural responses in all leave-one out folds and resulted in a significant linear prediction of the held out neural responses based on the linear fit (label permutation test, p<0.05), the unit was defined as a model neuron. Note that this search was carried out only in the DCNN layers that significantly matched the neural face-space (2 layers in each set, see **Figure 2A-C**). Finally, we applied a cluster correction to further ensure the received model neurons could not be detected by chance: we repeated the same search procedure 1000 times, each time shuffling the face labels of the neural responses. This resulted in a distribution of 1000 model neurons counts detected under shuffled data. Only if the original number of detected model neurons in a given layer exceeded the 95^th^ percentile of the count distribution derived from shuffled labels, we qualified the model neurons found in the layer as significant. For set 1, we found 7 and 2 model neurons in layer pool5 and layer conv5-3, respectively. For set 2, we found a single model neuron that did not survive cluster correction.

### Receptive field visualization of model neurons

We used two techniques to visualize the receptive field of detected model neurons. The first technique is commonly referred to as deconvolution and was established by Zeiler and Fergus (Güçlü & van Gerven, 2015; Zeiler & Fergus, 2014). Their method allows to feedforward an input image through a network, and then propagate it backwards to the image space given solely its representation in a unit of interest (i.e. setting all other units in the layer of the target unit to zero). The resulting back-propagated image thus unveils the receptive field of a defined target unit in the network as casted on the original input image – an approximation of the weighted fragment in the original image that gave rise to the target unit’s response. In practice, we based our implementation of the deconvolution technique of publicly available TensorFlow implementation (Bhagyesh & Falak, 2017).

The second technique that we used is commonly termed activation maximization (Erhan, Bengio, Courville, & Vincent, 2009). Here, we sought to *iteratively* alter the input image in a way that, once fed into the network again, would maximally increase the response of a target model neuron. In our implementation, we applied the decovolution technique iteratively (300 iterations): on each iteration we estimated the receptive field of the target unit using decovolution and added the deconvolved image, multiplied by a learning rate of 200, to the initial input image of that iteration. The resultant image was then used as the input image for the subsequent iteration. The visualizations presented in **Figure 4** and **Figure S6** are the delta between the rgb channels of the deconvolved image at the last iteration and the original image.

### ROI analysis

To investigate potential differences in the functional role of face patches along the posterior-anterior axis by means of comparison to DCNN layers (**Figure S5**), we defined two anatomical clusters in set 1 and 2 (set 3 did not include a sufficient amount of contacts to perform an ROI analysis). Face contacts assigned to the posterior cluster were defined as contacts located in up to 2 mm proximity to the left hemisphere inferior occipital gyrus. By contrast, face contacts assigned to the anterior cluster were defined as contacts located in up to 2 mm proximity to the left hemisphere fusiform gyrus. The subsequent analysis of the match between representational distances in the DCNN and in neural data was identical to that performed on the entire set of face contacts (**Figure 2**).

## Acknowledgments

This study has received funding from the CIFAR-Azrieli program of mind, brain and consciousness (Grant no.: 7129380101) and from the European Research Council (ERC) under the European Union’s Horizon 2020 research and innovation programme (Grant no. 788535). We would like thank Dr. Erez Simony for his helpful inputs in the process of preparing this work.

